# Transcriptome analysis of Pv11 cells infers the mechanism of desiccation tolerance and recovery

**DOI:** 10.1101/368175

**Authors:** Takahiro G Yamada, Yoshitaka Suetsugu, Ruslan Deviatiiarov, Oleg Gusev, Richard Cornette, Alexander Nesmelov, Noriko Hiroi, Takahiro Kikawada, Akira Funahashi

## Abstract

The larvae of the African midge, *Polypedilum vanderplanki*, can enter an ametabolic state called anhydrobiosis to conquer fatal desiccation stress. The Pv11 cell line, derived from embryos of the midge, shows desiccation tolerance by pretreatment with trehalose before desiccation; they can resume proliferation after rehydration. To address the underlying molecular mechanisms, we desiccated Pv11 cells after pretreatment with the medium containing trehalose and induced proliferation by rehydration. We collected the cells at each before and after desiccation and rehydration step and performed CAGE-seq of mRNA of those cells. By analysing differentially expressed genes (DEGs) among the results of CAGE-seq, we detected 384 DEGs after trehalose treatment and 14 DEGs after rehydration. Hierarchical clustering of the identified DEGs indicated that rehydration returns their expression pattern to that in the control culture state. DEGs involved in various stress responses, detoxification of harmful chemicals, and regulation of oxidoreduction were upregulated by trehalose treatment. DEGs for rehydration supported that DNA repair is one of the potential mechanisms involves recovery. This study provided initial insight into the molecular mechanisms underlying the extreme desiccation tolerance of Pv11 cells with a potential for proliferation following rehydration.

## Introduction

Desiccation stress, the loss of essential water, can be fatal. To tolerate desiccation stress, various organisms, such as rotifers, tardigrades, nematodes, plants, and larvae of the African midge *Polypedilum vanderplanki*, enter an ametabolic state called anhydrobiosis^1,2^ and survive even if more than 99% of body water is lost^3^. According to the water replacement hypothesis, a compatible solute, such as trehalose, protects phospholipid membranes and intracellular biological molecules and ensures their preservation under desiccation^3,4^. Consecutive desiccation can lead to serious oxidative stress. For example, in the moss *Fontinalis antipyretica* an increase in the production of reactive oxygen species (ROS) is associated with dehydration^5,6^. Protein oxidation in the dehydrated cells of yeast *Saccharomyces cerevisiae* is 10 times that of hydrated cells^5,7^. Thioredoxins (TRXs) remove harmful ROS and protect cells from ROS-induced damage^5,7^. The genome of *P. vanderplanki* has a paralogous gene cluster for TRXs^8^. These TRXs are upregulated by dehydration, and *P. vanderplanki* becomes tolerant to ROS-induced damage^8^. Upon rehydration, the anhydrobiotes return to active life.

In 2010, the Pv11 cell line was established as an embryonic cell culture from *P. vanderplanki*^9^. Desiccation tolerance of Pv11 cells is induced by treatment with culture medium containing 600 mM trehalose for 48 h. Even after dehydration in a desiccator (<10% relative humidity) for 12 days and rehydration for 1 h, the trehalose-treated Pv11 cells were able to resume proliferation^10^, whereas the other insect cell lines (Sf9, BmN-4, AeAl-2, AnCu-35, and S2) did not proliferate after rehydration. Pv11 cells are considered the only desiccation-tolerant insect cell line able to restore the regular cell cycle after rehydration, but it is unclear whether rehydrated Pv11 cells genuinely return to the same state. Analysis of gene expression patterns of Pv11 cells during the treatment, desiccation, and rehydration may help to elucidate which genes are required to avoid cell death and return Pv11 cells from anhydrobiosis to the normal state.

In this study, we performed high-throughput CAGE-seq of mRNA and differentially expressed gene (DEG) analysis in trehalose-treated, desiccated, and rehydrated Pv11 cells, followed by gene ontology (GO) analysis of the identified DEGs. To the best of our knowledge, this is the first report of comprehensive DEG analysis using CAGE-seq data for the Pv11 cells that infers a putative mechanism of their avoidance of cell death and recovery from anhydrobiosis.

## Results and Discussion

### Gene expression analysis by CAGE-seq

To detect genes potentially related to desiccation tolerance and successful recovery after rehydration, we analysed DEGs in control Pv11 cells (T0) and at different stages of anhydrobiosis: trehalose treatment for 48 h (T48), desiccation for 8 h or 10 days (D8 and D10d), and rehydration of D10d cells for 3 or 24 h (R3, R24). We compared each pair of samples (false discovery rate [FDR]<0.05; Table 1, Supplementary Data 1) and visualized the overall and differential expression of genes for each pair of samples as M-A plots (Supplementary Fig. S1). All of the median M values of non-DEGs were closer to 0 than those of DEGs; no DEGs were detected between D8 and D10d because metabolism of Pv11 cells completely stops under both these conditions. Samples were clustered using a hierarchical clustering algorithm (Fig. 1). The patterns of mRNA expression in the completely desiccated samples D8 and D10d were the most different from that of the normally cultured samples (T0). The expression patterns of R3 and R24 gradually approached that of T0.

**Figure 1.**
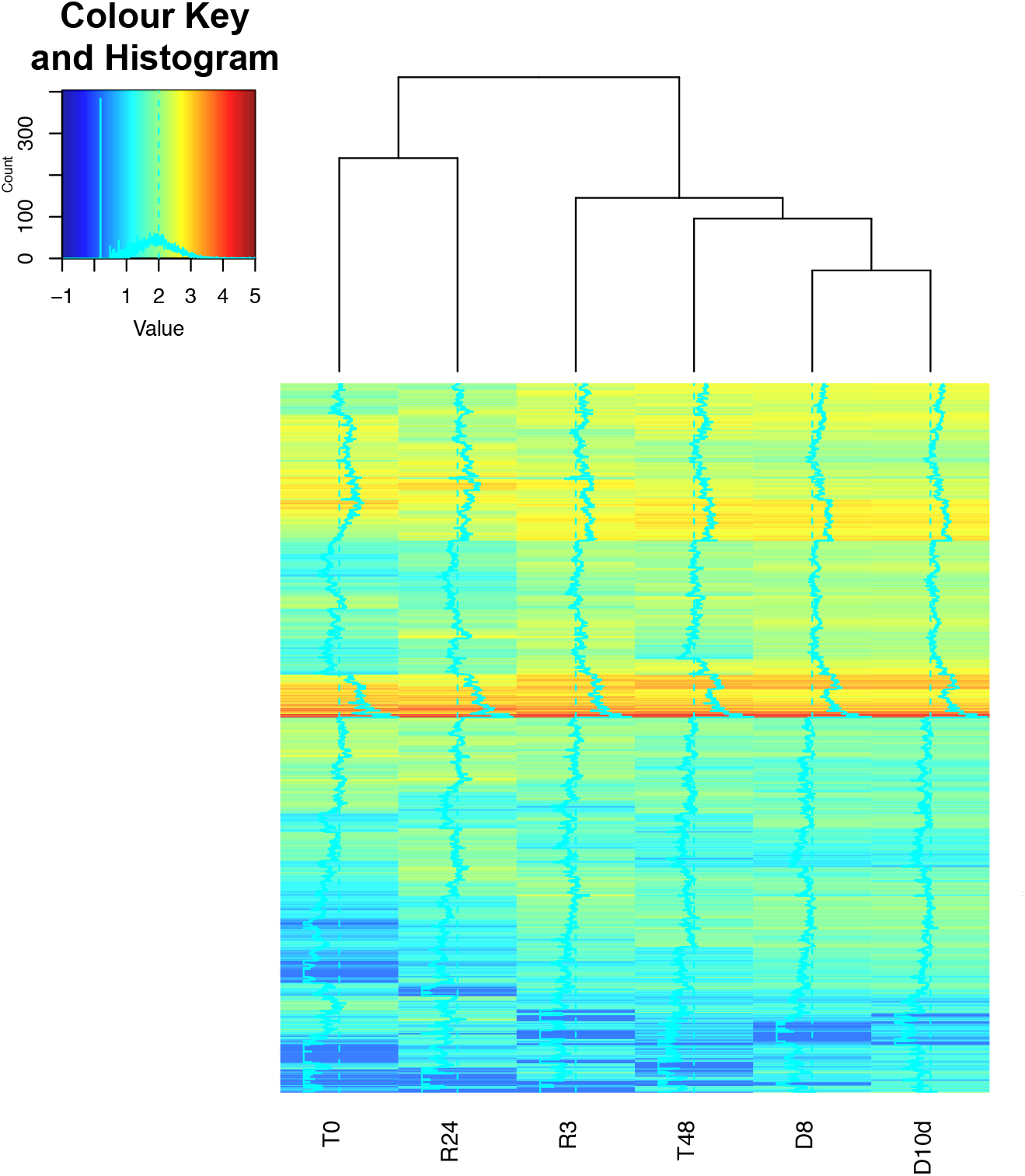
Hierarchical clustering based on tags per million (TPM) for each sample (distance: Euclidean). In the Colour Key and Histogram inset, the horizontal axis shows *log*_10_(*TPM* + *minimum TPM excluding* 0) and the vertical axis shows the number of DEGs. Genes were selected when they were detected as DEGs in at least one pair of samples.

**Table 1.**
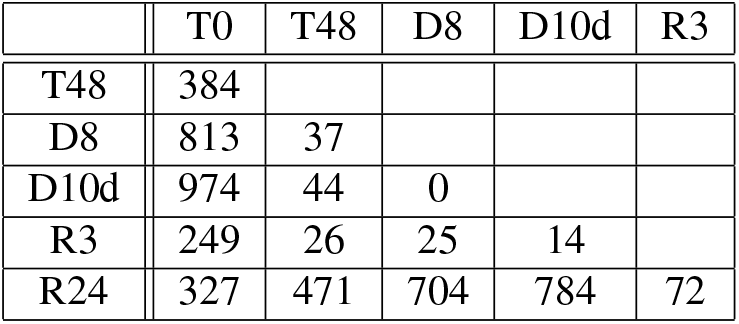
The numbers of differentially expressed genes. Each sample was analysed in biological triplicate, and the gene was considered as a DEG at FDR < 0.05 (Benjamini–Hochberg method). We analysed DEGs in control Pv11 cells (T0) and at different stages of anhydrobiosis: trehalose treatment for 48 h (T48), desiccation for 8 h or 10 days (D8 and D10d), and rehydration of D10d cells for 3 or 24 h (R3, R24)

|  | T0 | T48 | D8 | D10d | R3 |
| --- | --- | --- | --- | --- | --- |
| T48 | 384 |  |  |  |  |
| D8 | 813 | 37 |  |  |  |
| D10d | 974 | 44 | 0 |  |  |
| R3 | 249 | 26 | 25 | 14 |  |
| R24 | 327 | 471 | 704 | 784 | 72 |

### Gene Ontology analysis of DEGs

We expected that DEGs between T0 and T48, and D10d and R3 would be genes functionally important for desiccation tolerance and recovery from anhydrobiosis, respectively. We detected 299 upregulated and 85 downregulated DEGs between T0 and T48 (Supplementary Data 1). No enriched GOs for DEGs between T0 and T48 were detected (FDR<0.05, Supplementary Data 2). Therefore, we analysed the frequency of GOs among these DEGs. The first and second most frequent GOs were GO:0008152 (metabolic process) and GO:0055114 (oxidation-reduction process) in the biological process category (Fig. 2a), GO:0005524 (ATP-binding) and GO:0008270 (zinc ion binding) in the molecular function category (Fig. 2b), and GO:0005634 (nucleus) and GO:0016021 (integral component of membrane) in the cellular component category (Fig. 2c). Because metabolism and oxidoreduction are crucial for dehydration^3,8^, the genes with these GOs may be important for the induction of the desiccation tolerance mechanism.

**Figure 2.**
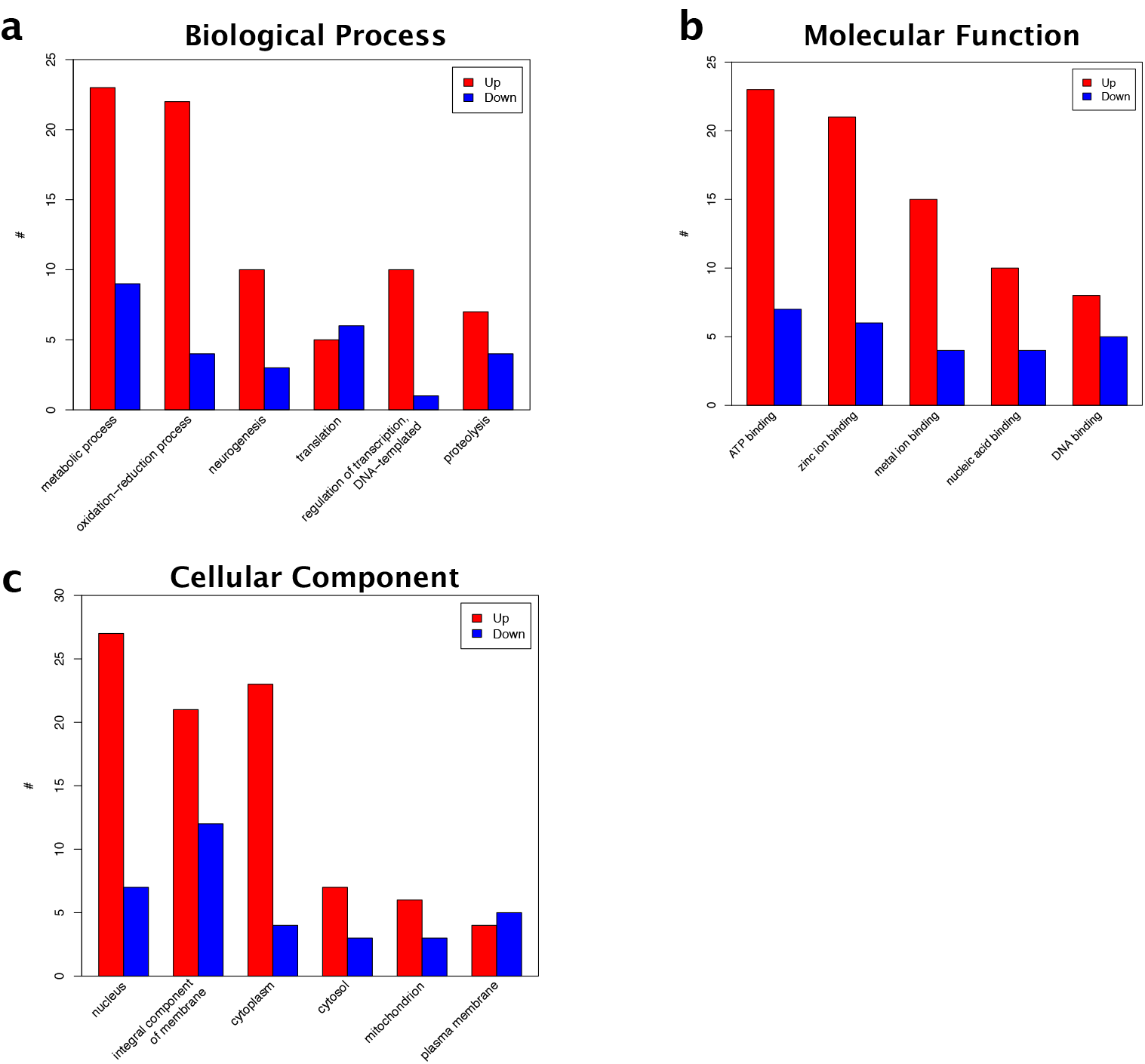
The frequency of Gene Ontology items in the annotations of the differentially expressed genes between T0 and T48.

In recovering cells, we detected 7 upregulated and 7 downregulated genes (Supplementary Table S1, Supplementary Data 1). GO analysis of these 14 genes identified 8 most significantly enriched GOs (Supplementary Table S2). The gene Pv.01867 was annotated by GO:0048763 (calcium-induced calcium release activity) and GO:0032237 (activation of store-operated calcium channel activity). Pv.04558 was annotated by GO:0030478 (actin cap) and GO:0070252 (actin-mediated cell contraction). Pv.07646 was annotated by GO:0036055 (protein-succinyllysine desuccinylase activity), GO:0036049 (peptidyl-lysine desuccinylation), GO:0036054 (protein-malonyllysine demalonylase activity), and GO:0036047 (peptidyl-lysine demalonylation). Pv.07646 was most significant among DEGs between D10d and R3; it was upregulated specifically in the R3 stage in some biological replicates (35 times vs. D10d and 15.5 times vs. R24; Supplementary Fig. S2a). RT-qPCR confirmed significant up-regulation of Pv.07646 (Supplementary Fig. S2b, Welch’s t test: *p*-value < 0.05). The two other genes were not specifically upregulated in R3 (Pv.01867: 4.93 times for R3 vs. D10d and 0.99 for R3 vs. R24; Pv.04558: 0.12 for R3 vs. D10d and 0.11 for R3 vs. R24; Supplementary Fig. S3).

### Analysis of desiccation tolerance mechanism

#### Hierarchical clustering of DEGs between T0 and T48

To reveal the biological function of the genes that participate in the induction of desiccation tolerance in Pv11 cells, we homed in on the DEGs between T0 and T48 with GO:0008152 (metabolic process; 32 DEGs) and GO:0055114 (oxidation-reduction process; 26 DEGs). To categorize these genes, we performed hierarchical clustering based on the data for each treatment. The expression patterns of GO:0008152 DEGs were classified into 4 clusters (Fig. 3a, Table 2). During desiccation after trehalose treatment, some DEGs were expressed consistently (Cluster1, Supplementary Fig. S4), but many were downregulated (Clusters 2–4, Supplementary Fig. S4). The expression patterns of DEGs with GO:0055114 were classified into 3 clusters (Fig. 3b, Table 3). During desiccation, the expression of DEGs in clusters 1 and 3 was maintained or upregulated, whereas DEGs in cluster 2 were drastically downregulated (Supplementary Fig. S5). The differences in the expression patterns of DEGs with the same GO during trehalose treatment, desiccation, and rehydration suggest different roles of these genes that enable Pv11 cells to acquire desiccation tolerance.

**Table 2.** Differentially expressed genes between T0 and T48 with GO:0008152 (metabolic process).“PREDICTED” is the status of initial RefSeq record. This status means the annotation is only predicted by automated BLAST analysis and is not validated by experiment.

| Cluster Num. | Pv Gene ID | Gene Identifier | BLAST First Hit Gene | E-value |
| --- | --- | --- | --- | --- |
| Cluster 1 | Pv.00012 | 755889065 | PREDICTED: uncharacterized protein LOC101899881 isoform X2 | 3.5155E-21 |
|  | Pv.00503 | 170031381 | conserved hypothetical protein | 0.0 |
|  | Pv.01170 | 195109112 | GI23236 | 0.0 |
|  | Pv.01614 | 557775558 | PREDICTED: tubulin beta-1 chain | 0.0 |
|  | Pv.01749 | 158296880 | AGAP008252-PA | 0.0 |
|  | Pv.01933 | 755849759 | PREDICTED: GTP-binding protein 1-like | 0.0 |
|  | Pv.03555 | 94468868 | NMD protein | 0.0 |
|  | Pv.05750 | 347969792 | AGAP003372-PB | 8.73E-102 |
|  | Pv.06865 | 157120582 | AAEL009068-PA | 0.0 |
|  | Pv.07614 | 568250312 | transport and golgi family organization | 2.44E-75 |
|  | Pv.07843 | 668455678 | hypothetical protein ZHAS_00011763 | 4.90E-162 |
|  | Pv.08737 | 170064729 | ATP-dependent RNA helicase A | 0.0 |
|  | Pv.11162 | 668458368 | AGAP004610-PA-like protein | 0.0 |
|  | Pv.12163 | 568253176 | DEAD box ATP-dependent RNA helicase | 3.32E-146 |
|  | Pv.13802 | 170049985 | conserved hypothetical protein | 2.60E-86 |
|  | Pv.15605 | 157107882 | AAEL014906-PA, partial | 2.81E-76 |
|  | Pv.17087 | 817062658 | PREDICTED: E3 ubiquitin-protein ligase rnf146-like | 1.19E-32 |
| Cluster 2 | Pv.02231 | 751789335 | PREDICTED: 5-oxoprolinase | 0.0 |
|  | Pv.06995 | 227343489 | glutathione S-transferase | 5.64E-87 |
|  | Pv.10576 | 347969074 | AGAP003018-PA | 0.0 |
|  | Pv.12167 | 157110933 | AAEL000854-PA | 9.32E-141 |
| Cluster 3 | Pv.02391 | 749772182 | PREDICTED: abhydrolase domain-containing protein 4 isoform X1 | 9.38E-112 |
|  | Pv.06862 | 568250143 | Elongator complex protein 3 | 0.0 |
|  | Pv.08047 | 332373216 | unknown | 8.53E-36 |
|  | Pv.11397 | 170054183 | UDP-glucuronosyltransferase 2C1 | 8.03E-30 |
|  | Pv.12035 | 157117142 | AAEL007889-PA | 0.0 |
|  | Pv.13081 | 157136208 | AAEL003433-PA | 0.0 |
|  | Pv.13347 | 557763449 | PREDICTED: E3 ubiquitin-protein ligase Topors isoform X2 | 3.02E-56 |
|  | Pv.16072 | 668446285 | AGAP008326-PA-like protein | 1.05E-145 |
| Cluster 3 | Pv.04950 | 568250792 | carboxylesterase, beta esterase | 1.67E-138 |
|  | Pv.06833 | 589278470 | hypothetical protein TREMEDRAFT_39272 | 7.49E-26 |
|  | Pv.13234 | 668450732 | hypothetical protein ZHAS_00007358 | 1.72E-28 |

**Table 3.**
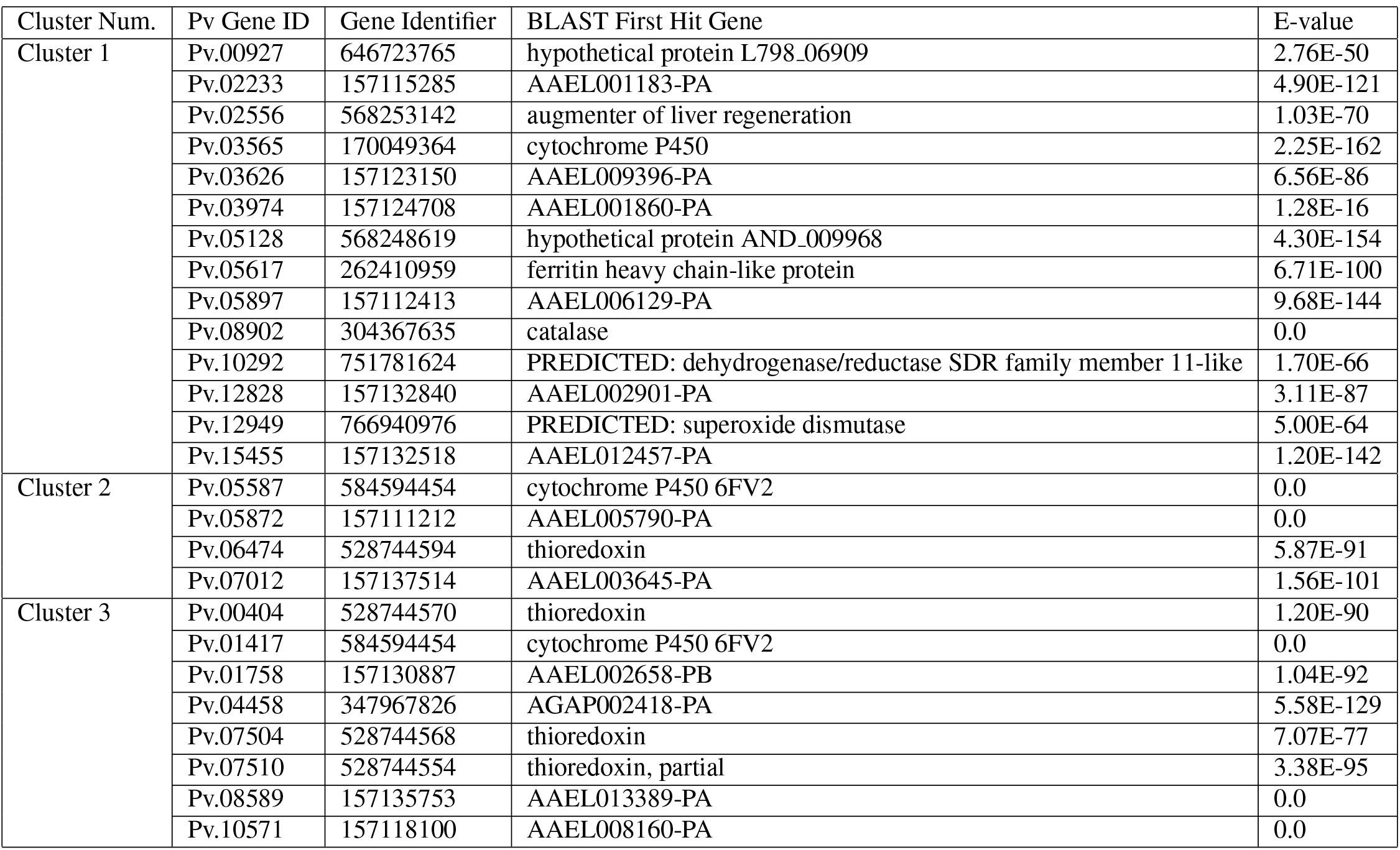
Differentially expressed genes between T0 and T48 with GO:0055114(oxidation-reduction process).

**Figure 3.**
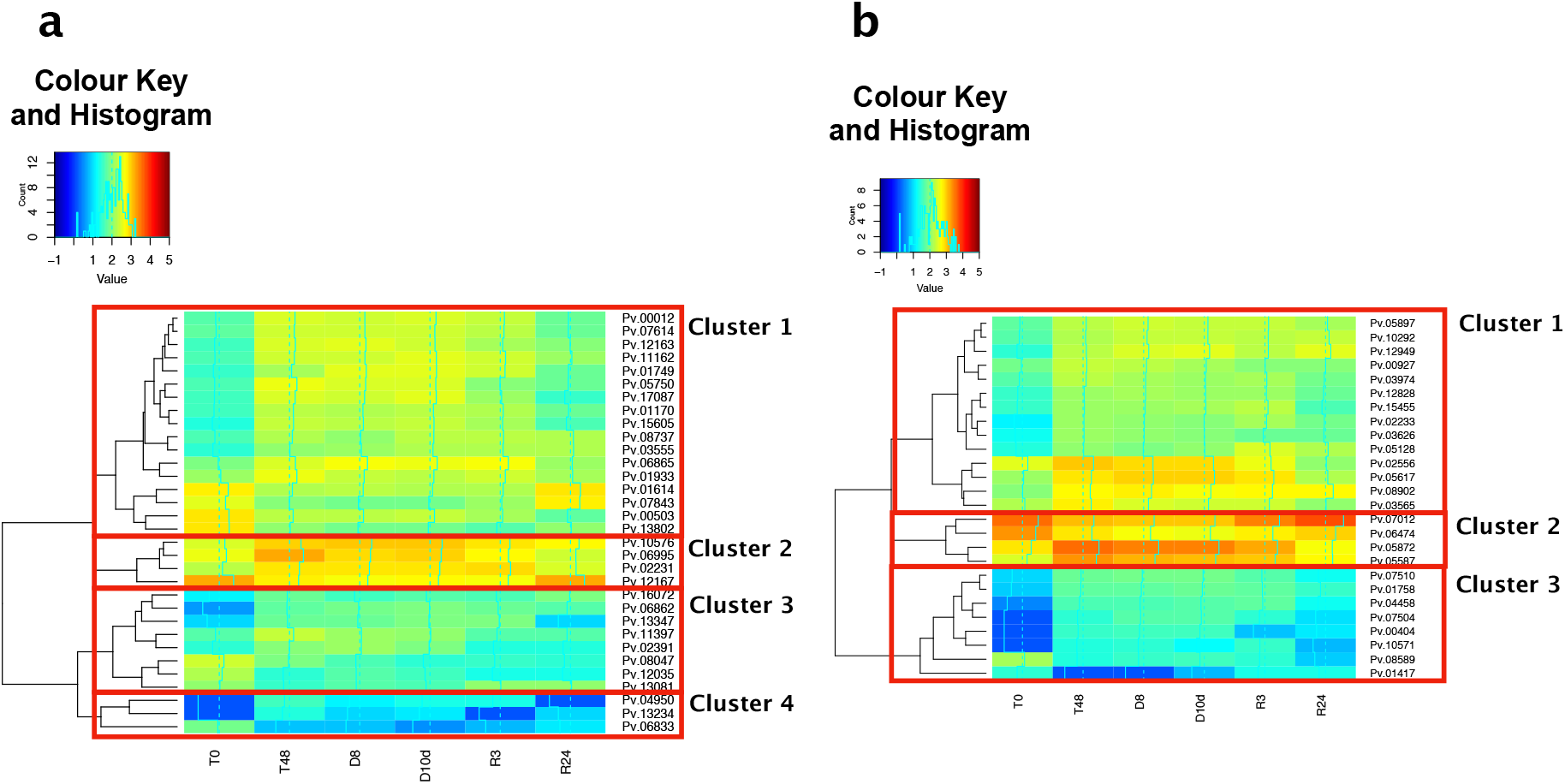
Hierarchical clustering of differentially expressed genes (DEGs) with (a) GO:0008152 (metabolic process) and (b) GO:0055114 (oxidation-reduction process). In the Colour Key and Histogram insets, the horizontal axes show *log*_10_(*TPM* + *minimum TPM excluding* 0) and the vertical axes show the number of DEGs.

#### Various protection systems in the induction of desiccation tolerance

To clarify the contributions of DEGs to the induction of desiccation tolerance, we focused on the functions of genes with GO:0008152 and GO:0055114 identified by BLASTX searches. Pv.03555 (GO:0008152) has the highest similarity to the nonsense-mediated mRNA decay (NMD) protein (Supplementary Fig. S6), which distinguishes mRNA containing a premature stop codon and induces its degradation^11^. Pv.06995 (GO:0008152) has the highest similarity to glutathione S-transferase (GST) (Supplementary Fig. S7). GST is critical for the protection against lipid peroxidation and for detoxification of oxidized lipids^12^. In *Arabidopsis thaliana*, upregulation of the gene encoding GST protects from drought stress^13^. Pv.03555 and Pv.06995 were significantly upregulated in T48 in comparison with T0, indicating that the expression of various stress response genes was induced by trehalose treatment in preparation for stress caused by dehydration. Pv.11397 (GO:0008152) has the highest similarity to UDP-glucuronosyltransferase (UGT) (Supplementary Fig. S8), which catalyses the glucuronidation of toxic chemicals such as bilirubin, phenols, amines, and carboxylic acids^14^. Therefore, Pv.11397 is likely to contribute to the detoxification of some harmful chemicals during induction of desiccation tolerance. In *Caenorhabditis elegans*, UGT is upregulated 6.5 fold after dehydration^15^. Pv.04950 (GO:0008152) has the highest similarity to carboxylesterase (Supplementary Fig. S9). Carboxylesterases catalyse the hydrolysis of xenobiotics with ester and amide groups^16^. Pv.11397 and Pv.04950 were significantly upregulated in T48 in comparison with T0. On the basis of the functions of these genes, we assume that their upregulation allows removal of harmful chemicals.

Many GO:0008152 genes were downregulated after dehydration, likely because Pv11 cells enter an ametabolic state after dehydration and metabolism-related genes may become unessential. On the other hand, the expression of GO:0008152 DEGs in Cluster 1 (Fig. 3) was maintained during dehydration. These DEGs included Pv.03555, which has the highest similarity to the NMD protein (as described above); Pv.08737, which has the highest similarity to ATP-dependent RNA helicase (Supplementary Fig. S10); and Pv.12163, which has the highest similarity to DEAD box ATP-dependent RNA helicase (Supplementary Fig. S11). Both NMD protein and RNA helicases act in the clearance of failed mRNA^17^. Therefore, the corresponding proteins of *P. vanderplanki* likely allow efficient elimination of imperfect mRNA that may result from strongly oxidizing conditions during dehydration.

Pv.00404, Pv.06474, Pv.07504, and Pv.07510 (oxidation-reduction process) have the highest similarity to thioredoxins (TRXs) and have been reported as PvTrx1–1, PvTrx3, PvTrx7, and PvTrx10, respectively^8^. These genes are upregulated after dehydration in the larvae of *P. vanderplanki*^8^. We found that these genes were significantly upregulated in T48 in comparison with T0. These findings indicate that oxidoreduction is part of the mechanism for the induction of desiccation tolerance and show an analogy between Pv11 cells and larvae of *P. vanderplanki*.

### Analysis of the mechanisms of recovery from anhydrobiosis

#### Domain structure analysis of the Pv.07646 gene

The alignment of Pv.07646 and its putative homolog SNF histone linker PHD ring helicase is shown in Supplementary Figure S12. Histone Lys residues are succinylated and malonylated^18^. To recognize succinylated or malonylated Lys residues, the enzyme involved in desuccinylation or demalonylation needs to bind both the histone and DNA. Pv.07646 satisfies these requirements because it has a RING domain, which may be involved in protein–protein interactions, and a nucleotide-binding site. However, Pv.07646 has no sequence similarity to the SIRT5 region, which is responsible for desuccinylation and demalonylation, suggesting that Pv.07646 may not directly catalyse these reactions. Pv.07646 has the highest similarity to the DNA-repair protein RAD16. The alignment analysis showed that Pv.07646 has sequences homologous to the HepA domain and the ATP-binding site of RAD16 (E-values, 0.0; Supplementary Fig. S13, Supplementary Table S1). Rad16 forms a complex with Rad7 and binds DNA in an ATP-dependent manner^19^. When the HepA region of Rad16 is deactivated and DNA is damaged by UV radiation, the survival rate of *Escherichia coli* is significantly decreased^20^. These results suggest that Pv.07646 may act in DNA repair.

#### DNA repair system in anhydrobiosis

We identified three DEGs that were returned as hits in BLASTX searches and whose names included “DNA repair” (Table 4). Pv.04176 has high similarity to the RAD51 (Supplementary Fig. S14); Pv.04176 was significantly upregulated in D10d in comparison with T0. RAD51 forms a complex with single-stranded DNA (ssDNA) in homologous recombination repair^21^. The RAD51–ssDNA filament is then stretched to as much as half the length of B-form double-stranded DNA. This stretching of the DNA is essential for accomplishing fast and efficient homology search^22^. After that, RAD51 facilitates the formation of a physical connection between the invading DNA substrate and homologous duplex DNA template. Finally, RAD51 is released from the filaments to expose the 3’-OH for DNA repair^21^. Therefore, Pv.04176 may function in the recognition, searching, and protection of broken DNA in homologous recombination repair. Pv.16797 is highly similar to XRCC1 (Supplementary Fig. S15); Pv.16797 was gradually upregulated by dehydration and rehydration. XRCC1 is a critical scaffold protein that interacts with, and in some cases stabilizes and/or stimulates, several enzymes of single-strand break repair^23^. Thus, Pv.16797 may function as a scaffold protein during dehydration and this function may precede DNA repair. Pv.10801 shares sequence similarity with XPA (Supplementary Fig. S16); Pv.10801 was significantly upregulated in R24 in comparison with D10d. The XPA protein assists in the disassembly of chromatin to make it accessible for incision nucleases^24^. Therefore, Pv.10801 may disassemble chromatin prior to DNA repair.

**Table 4.** The genes of *P. vanderplanki* that were the hits in BLASTX searches and whose names included “DNA repair”.

| Pv Gene Id | Gene Identifier | BLAST Hit Gene Name | E.value | Significant Upregulation in dehydrated samples (D8 or D10d) compared with other samples (FDR < 0.05) | Significant Upregulation in rehydrated samples (R3 or R24) compared with other samples (FDR < 0.05) |
| --- | --- | --- | --- | --- | --- |
| Pv.10243 | 557752841 | alkylated DNA repair protein alkB homolog 1 | 7.08E-65 | - | - |
| Pv.17500 | 170041242 | DNA repair and recombination protein RAD54 | 0.0 | - | - |
| Pv.10801 | 557761125 | DNA repair protein complementing XP-A cells homolog | 9.13E-78 | - | D10dvsR24 |
| Pv.03629 | 91087279 | DNA repair protein complementing XP-C cells homolog | 1.69E-36 | - | - |
| Pv.03630 | 751778114 | DNA repair protein complementing XP-C cells homolog | 2.05E-146 | - | - |
| Pv.07179 | 170035857 | DNA repair protein RAD50 | 3.36E-34 | - | - |
| Pv.02239 | 170035857 | DNA repair protein RAD50 | 0.0 | - | - |
| Pv.07016 | 170035857 | DNA repair protein RAD50 | 1.59E-58 | - | - |
| Pv.13318 | 498925123 | DNA repair protein RAD50 | 1.41E-53 | - | - |
| Pv.04176 | 304367643 | DNA repair protein RAD51 | 0.0 | T0vsD10d | - |
| Pv.04143 | 170048628 | DNA repair protein RAD62 | 3.47E-38 | - | - |
| Pv.16797 | 498935604 | DNA repair protein XRCC1 isoform X2 | 1.07E-72 | T0vsD8 and T0vsD10d | T0vsR3 |
| Pv.11679 | 568260085 | DNA repair/transcription protein met18/mms19 | 3.56E-88 | - | - |
| Pv.14185 | 170041817 | DNA repair/transcription protein met18/mms19 | 3.34E-96 | - | - |

### Suggested model

Treatment with medium containing sufficient trehalose is essential for Pv11 cells to induce desiccation tolerance; Pv11 cells treated with the culture medium containing 443 mM trehalose did not survive, but with an increase to 600 mM, the survival rate increased to 16%^10^. Trehalose is a compatible solute and protects the cell membrane and intracellular biological molecules^3,4^. The detection of 384 DEGs between T0 and T48 indicates that trehalose not only directly protects biological molecules but also regulates many genes for desiccation tolerance in Pv11 cells. The accumulation of trehalose in *P. vanderplanki* larvae starts just after the onset of desiccation and water content is steady at that time^25^. We assumed that treatment of Pv11 cells with trehalose corresponds to this time span and various genes needed to acquire desiccation tolerance are upregulated in *P. vanderplanki*.

Various stress response genes (Pv.03555 and Pv.06695), genes for proteins that eliminate harmful chemicals (Pv.11397 and Pv.04950), and TRX genes (Pv.00404, Pv.06747, Pv.07504, and Pv.07510) were significantly upregulated in T48 in comparison with T0. The known biological functions of similar genes suggest that all these genes likely have a role in protecting cells from the harmful effects of desiccation.

On the basis of the similarity of domain structure between Pv.07646 and Rad16, we suggest that Pv.07646 could be involved in DNA repair during rehydration. DNA repair is required for survival in plants and prokaryotes after anhydrobiosis^26,27^. Tardigrades also recover if DNA damage in the desiccated state is low^28^. The differential expression of DNA repair gene homologs in Pv11 cells during dehydration and at R3 suggests that DNA damage accumulated during anhydrobiosis must be repaired before cells can resume normal growth. The RAD51 homolog Pv.04176 may protect DNA damaged during dehydration, and the XRCC1 homolog Pv.16797 may scaffold DNA during dehydration and rehydration. Then, as cells are rehydrated, the XPA homolog Pv.10801 may interact with chromatin and possibly the RAD16 homolog Pv.07646, detect DNA damage and prepare DNA for repair.

## Methods

### CAGE RNA preparation, Illumina Hiseq-2500 sequencing, and data preprocessing

Pv11 cells were maintained in IPL-41 medium supplemented with 10% FBS (T0) and were incubated in 600 mM trehalose solution containing 10% IPL-41 medium for 48 h (T48). The cells were desiccated in a desiccator (relative humidity, <10%) for 8 h (D8) or 10 days (D10d). After D10d, the samples were rehydrated with IPL-41 containing 10% FBS for 3 h (R3) or 24 h (R24). Samples were prepared in biological triplicate. Total RNA was extracted with a RNAiso-Plus kit (TaKaRa, Shiga, Japan) and purified with a NucleoSpin^®^RNA kit (Macherey-Nagel, Düren, Germany). RNA quality was assessed with a Bioanalyzer (Agilent Technologies, Santa Clara, CA, USA), and RNA was quantified with a Qubit 2.0 fluorometer (Thermo Fisher, Waltham, MA, USA). First-strand cDNAs were synthesized on the 5’-ends of capped RNAs attached to CAGE bar code tags. A CAGE library was prepared as described previously^29^. CAGE tags were sequenced with HiSeq-2500 (Illumina, San Diego, CA, USA). The reads were preprocessed by removing (a) the first 3 bases from the 5’-end considering it could be capped, (b) the first 7 bases from the 3’-end if they were contiguous adenine residues because it could be a poly-A structure; and (c) reads containing more than 2 N bases because mapping such reads to the genome could result in false positives (Supplementary Table S3). The remaining reads were mapped to the genome of *P. vanderplanki* (midgeBase: http://150.26.71.110/midgebase/) using the qAlign function in the QuasR package in Bioconductor^30^. The mapped reads were counted using the qCount function in the QuasR package.

### DEG analysis

The read count data of each sample were revised using the tags per million (TPM) and then using the iterated DEG Eliminated Strategy (iDEGES) method (TMM-(edgeR-TMM)_3_)^31^. Using the revised data, DEGs were detected by edgeR^32^ when false discovery rate (FDR) calculated by the Benjamini–Hochberg method^33^ was less than 0.05. The differential expression of genes for each pair of samples was plotted as an M-A plot^34^. The horizontal axis of the M-A plot indicates the average expression of a gene across two groups (samples G1 and G2).

### Heat map and hierarchical clustering

The TPMs of DEGs among all samples were visualized as a heat map (CRAN: heatmap.2;^35^). The similarity of the expression pattern between samples or genes was analysed by hierarchical clustering (CRAN: hclust;^36^). The distance metric and linkage method were Euclidean distance and complete linkage method, respectively.

### Gene Ontology annotation and analysis by Fisher’s exact test

To assess the similarity of primary structures, BLAST searches of the non-redundant database were performed using Blast2GO^37^ with the BLASTX option and an E-value threshold of 1.0E-05. GO mapping and annotation of *P. vanderplanki* genes were performed on the basis of the BLASTX results with the thresholds of CutOff and GO level weighting set at 30 and 5, respectively (Supplementary Data 3). GO analysis was performed using Fisher’s exact test to detect enriched GO terms in the annotations. The probability of detecting a GO term X in a set of DEGs by chance was calculated as

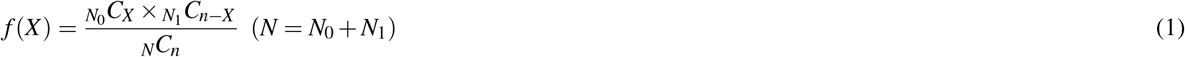

where *N* is the number of all GO terms, *N*_0_ is the number of noted GO terms, and *n* is the total number of all GO terms in the set of DEGs. The *p*-value to detect X by chance was calculated using the fisher.test function in R^36^ as

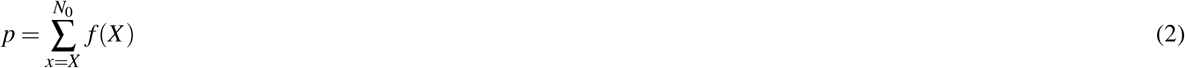

FDR was calculated by the Benjamini–Hochberg method using the *p*-value. GOs with FDR < 0.05 were selected. GOs with similar functions were gathered by REVIGO^38^. The method of semantic similarity calculation was set up with the Rel method^39^ with the default threshold of 0.7. The selected GOs were considered enriched.

### Domain structure analysis

The similarity of protein domains of gene products registered in the Conserved Domain Database (CDD)^40^ was evaluated by using a multiple sequence alignment viewer (MSAViewer:https://www.ncbi.nlm.nih.gov/projects/msaviewer/;^41^). Designations are as in Supplementary Figure S6.

### RT qPCR

D10d and R3 samples were prepared as in CAGE-seq data preparation. Cells were homogenized by ultrasound. Total RNA was extracted using a ReliaPrep RNA Tissue Miniprep System and quantified using a QuantiFluor ssDNA System (both from Promega Corporation, Madison, WI, USA). RNA was reverse transcribed into cDNA with a Transcriptor First Strand cDNA Synthesis Kit (Roche Applied Science, Mannheim, Germany) using an anchored oligo (dT) random hexamer primer (600 pmol/*μ*L) for 60 min at 60°C and 5 min at 85°C. cDNA was then immediately placed on ice, quantified using the QuantiFluor ssDNA System, and diluted with MilliQ water to contain 10 ng of cDNA based on the quantification result. qPCR was performed in a CFX96 Real-Time PCR Detection System (Bio-Rad, Hercules, CA, USA) for 30 s at 95°C and 41 cycles of 5 s at 95°C and 30 s at 62°C. Each qPCR mixture contained 10 *μ*L of SYBR Premix Ex Taq (Tli RNaseH Plus) (2×) (TaKaRa), 10 ng of cDNA, and 0.2 *μ*M each forward and reverse primers, supplemented with MilliQ water to 20 *μ*L. The primers used for Pv.07646 were 5’-TGAGACAAGACGAGCCAGATG-3’ (forward) and 5’-CCAAATGAGGAGCGGAATG-3’ (reverse). A control reaction without the template was also performed. Fold change of gene expression in R3 relative to D10d was calculated as 2^*C_t_* (*D*10*d*)−*C_t_* (*R*3)^, where *C_t_* is the threshold cycle.

## Data availability

CAGE-seq data of mRNA is available on request from the corresponding author. All other data generated or analysed during this study are included in this published article and its Supplementary Information files.

## Acknowledgements

We are grateful to Tomoe Shiratori for maintenance of Pv11 cells and to Yuki Sato-Kikuzato for assistance with library preparation. The research was funded by JSPS KAKENHI (Grant Numbers 25128714 and 17H01511) from the Agriculture, Forestry and Fisheries Research Council of the Ministry of Agriculture, Forestry and Fisheries of Japan, Russian Science Foundation Joint Research Groups Grant 17–44-07002, and Russian Science Foundation grant 17–44-00003. The manuscript has been carefully checked by Ryo J. Nakatani.

## Author contributions statement

T.K. and A.F. conceived and led the project. T.Y. analysed data and wrote the manuscript. O.G., R.C., R.D., and A.N. designed, prepared and sequenced CAGE-seq libraries. N.H. and T.K. provided biological expertise. A.F. gave technical advice on analyses. Y.S. adviced DEG analysis and developed the genome database of *Polypedilum vanderplanki* (midgeBase). All authors were involved in drafting or revising the content of the manuscript. All authors read and approved the manuscript.

## Additional information

### Competing interests

The authors declare no competing interests.

